# Diploid male production correlates with genetic diversity in the parasitoid wasp *Venturia canescens*: a genetic approach with new microsatellite markers

**DOI:** 10.1101/054866

**Authors:** Marie Collet, Chloé Vayssade, Alexandra Auguste, Laurence Mouton, Emmanuel Desouhant, Thibaut Malausa, Xavier Fauvergue

**Affiliations:** Université de Lyon, Université Claude Bernard, CNRS, Laboratoire de Biométrie et Biologie Evolutive UMR 5558, F-69622 Villeurbanne, France; INRA, Université Nice Sophia Antipolis, CNRS, UMR 1355-7254 Institut Sophia Agrobiotech, 06900 Sophia Antipolis, France

**Keywords:** microsatellite markers, sl-CSD, *Venturia canescens*, diploid males

## Abstract

Sex determination is ruled by haplodiploidy in Hymenoptera, with haploid males arising from unfertilized eggs and diploid females from fertilized eggs. However, diploid males with null fitness are produced under Complementary Sex Determination (CSD), when individuals are homozygous for this locus. Diploid males are expected to be more frequent in genetically eroded populations (such as islands and captive populations), as genetic diversity at the csd locus should be low. However, only a few studies have focused on the relation between population size, genetic diversity and the proportion of diploid males in the field. Here, we developed new microsatellites markers in order to assess and compare genetic diversity and diploid male proportion in populations from three distinct habitat types (mainland, island or captive), in the parasitoid wasp *Venturia canescens*. Eroded genetic diversity and higher diploid male proportion were found in island and captive populations, and habitat type had large effect on genetic diversity. Therefore, diploid male proportion reflects the decreasing genetic diversity in small and isolated populations. Thus, Hymenopteran populations can be at high extinction risk due to habitat destruction or fragmentation.

## Introduction

Sex determination takes on many different forms in animals (Bull, 1983). In insects of the order Hymenoptera, haplodiploidy is the rule: males develop from unfertilized eggs and are haploid whereas females develop from fertilized eggs and are diploid. In numerous species including sawflies, pollinating bees, social and solitary wasps, ants and parasitoids, the underpinning genetic mechanism of sex determination relies on the complementarity of alleles at a single gene (Heimpel and de Boer, 2008; Whiting, 1943). This gene-*csd*, for complementary sex determiner-has been described in the honeybee, and is probably responsible for sex determination in all species of the order Hymenoptera (Asplen et al., 2009; Beye et al., 2003; Schmieder et al., 2012). Under single locus complementary sex determination (sl-CSD), unfertilized eggs are hemizygous at the csd locus and develop normally into haploid males (Whiting, 1943). Among the fertilized eggs, only heterozygous individuals develop into females;homozygosity at the csd locus yields diploid males that are generally, but not always,unviable or sterile (Cowan and Stahlhut, 2004; Harpur et al., 2013). sl-CSD is the mechanism of sex determination for which empirical evidence prevails in Hymenoptera (Cook, 1993; van Wilgenburg et al., 2006), but other mechanisms such as multi-locus CSD or genomic imprinting also occur (Heimpel and de Boer, 2008).

At the population level, single-locus complementary sex determination causes higher sensibility to losses of genetic diversity. A decrease in allelic richness at the csd locus and a subsequent increase in homozygosity is expected to result in more diploid males and, in turn, lower population growth rate. Theoretical models predict that sl-CSD is at the core of a “Diploid Male Vortex”, a feedback between demographic and genetic processes that potentially drives bottlenecked populations toward extinction (Bompard et al., submitted; Fauvergue et al., 2015; Hein et al., 2009; Zayed and Packer, 2005).However, diploid males represent such a fitness cost that various individual behaviours have evolved to limit their production. Dispersal limits inbreeding and the consequent diploid male production (Ruf et al., 2011). Dispersal also allows the recruitment of new *csd* alleles, so that well-connected populations are expected to produce less diploid males (Hein et al., 2009). Some Hymenoptera with sl-CSD have undifferentiated populations beyond a 100 km scale (Estoup et al. 1996; Zimmermann et al. 2011) suggesting good dispersal abilities. Mate choice can also influence the production of diploid males with a reduction expected in case of inbreeding avoidance (Metzger et al., 2010a; Thiel et al., 2013; Whitehorn et al., 2009). Such behaviours may mitigate the diploid male vortex (Hein et al., 2009).

The central assumption of CSD population models is that small, bottlenecked or isolated populations have a low genetic diversity, including diversity at the *csd* locus, and consequently, a high proportion of diploid males. To date, however, empirical support to this assumption is scarce. Only rare field data on fire ants (Ross et al., 1993), bumblebees (Darvill et al., 2012, 2006) or paper wasps(Tsuchida et al., 2014) suggest a relation between population size, genetic diversity and the proportion of diploid males. These studies have all relied on genetic markers to estimate within-population genetic diversity, gene flow between distinct populations, mating structure and, in some cases, the proportion of diploid males.

Evidence for diploid males in natural populations mainly concerns social Hymenoptera. In these species, the dramatic bias in sex ratio toward females renders estimations of the proportion of diploid males very tedious. In parasitoid wasps, the proportion of diploid males was assessed in only three related species: *Cotesia glomerata* (Ruf et al., 2013), *C. rubecula* (de Boer et al., 2012) and *C. vestalis* (de Boer et al., 2015).Parasitoid populations often experience small population size, e.g. as a consequence of cyclic dynamics (Hassell, 2000) or biological control introductions (Hopper and Roush, 1993). Given the key role that parasitoid wasps play by controlling the populations of herbivorous insects in natural ecosystems and agrosystems (Shaw and Hochberg, 2001), the scarcity of field estimations on the occurrence of diploid males in the field is surprising.

*Venturia canescens* G. (Hymenoptera: Ichneumonidae) is a parasitoidwasp with sl-CSD(Beukeboom, 2001). This species is a model organism commonly used in laboratory studies on behaviour, physiology and life-history traits (Desouhant et al., 2005; Harvey et al., 2001; Pelosse et al., 2007). Two reproduction modes are known in this species (Schneider et al., 2002): thelytoky (asexual, parthenogenesis) and arrhenotoky (sexual reproduction under haplodiploidy). In arrhenotokous *V. canescens*, diploid males are fully viable and able to mate. However, they are sterile, and females withwhom they mateproduce only sons, similarly to virgin females (Fauvergue et al., 2015). The prevalence of diploid males in natural or captive *V. canescens* populationshas not been estimatedyet, and little is known about the genetic structure of natural populations. So far, the rare population genetic studies on *V. canescens* showed an absence of genetic structure at the geographic scale of the French Riviera (Mateo Leach et al., 2012; Schneider et al., 2002).

We developed the present study with a threefold objective: (i) develop microsatellite markers in order to assess male ploidy and conduct population genetic studies on arrhenotokous *V. canescens*; (ii) estimate the proportion of diploid males in several populations; (iii) estimate the genetic diversity of *V. canescens* populations under various conditions of population size and isolation. For this, we compared mainland, island and captive populations, with the expectation of a lower genetic diversity and a higher proportion of diploid males in captive populations compared to insular populations, and in insular populations compared to mainland populations.

## Materials and methods

### Sampling

Females *Venturia canescens* were collected during the summer 2010 in two natural populations located in Southern France near Nice and Valence (Table 1). To attract parasitoid females, open cups containing a mixture of host-rearing medium (semolina) and larvae of the host *Ephestia kuehniella* (Lepidoptera: Pyralidae) were hung in trees. Infested semolina is impregnated with host kairomones and is very attractive for *V. canescens* females (Corbet 1971; Metzger *et al.* 2008). Females visiting the traps to oviposit were collected, brought into the laboratory, allowed to lay eggs for two days on a host patch, and then killed and preserved in 96%ethanol. The reproductive mode of captured females was tested by checking the presence of males among offspring. Only arrhenotokous (sexual) females were used in the following analysis.

**Table 1.** Studied populations: locality, habitat type (mainland, island, captive populations), geographic coordinates, host plant (Car, Fig, Wal, Pom, Che, Pea, Haz, Cit and Oli being respectively Carob, Fig, Walnut, Pomegranate, Cherry, Peach, Hazelnut, Citrus and Olive trees), year of sampling and year of foundation between parentheses for captive populations, with the corresponding number of males or females sampled. See Figure 1 for a map of localities.

| Habitat type | Populations |  |  |  |  | Sampling |  |  |
| --- | --- | --- | --- | --- | --- | --- | --- | --- |
|  | Country | Locality | Name | Geographic coordinates | Host plant | Date | Number of males | Number of females |
| Mainland | France | Valence | Val10 | N 44° 58' 21"<br>E 4° 55' 39" | Che/Pea/Haz | 2010 | 0 | 37 |
|  |  | Nice | Nice10 | N 43° 41' 23"<br>E 7° 18' 6" | Car | 2010 | 0 | 44 |
|  |  | Nice | Nice11 | N 43° 41' 23"<br>E 7° 18' 6" | Car | 2011 | 190 | 0 |
|  |  | Nice | Nice13 | N 43° 41' 23"<br>E 7° 18' 6" | Car | 2013 | 90 | 21 |
|  |  | Solliès | Sol13 | N 43° 10' 58"<br>E 6° 2' 55 " | Fig/Wal/<br>Pom/Che | 2013 | 16 | 0 |
|  | Spain | Vila-Seca | VS13 | N 41° 7' 34"<br>E 1° 8' 07" | Car | 2013 | 58 | 2 |
|  |  | Vinyols | Vy13 | N 41° 6' 12"<br>E 1° 2' 26" | Car/Oli | 2013 | 33 | 2 |
| Island | France | Corsica | NA† |  |  | 2013 | 0 | 0 |
|  |  | Porquerolles | NA† |  |  | 2013 | 0 | 0 |
|  | Spain | Mallorca | Mlc12 | E 6° 13' 18"<br>E 2° 57' 53" | Car | 2012 | 21 | 0 |
|  |  | Mallorca | Mlc13 | N 39° 47' 58"<br>E 2° 57' 53" | Car | 2013 | 38 | 3 |
|  | Italy | Sicily | NA† |  |  | 2012 | 0 | 0 |
|  | Greece | Crete | Cre12 | N 35° 11' 39"<br>E 25° 2' 09" | Pom/Fig/Cit/Oli | 2012 | 0 | 1 |
|  |  | Crete | Cre13 | N 35° 11' 39"<br>E 25° 2' 09" | Pom/Fig/Cit/Oli | 2013 | 0 | 1 |
|  | Malta | Gozo | Goz12 | N 36° 3' 34"<br>E 14° 16' 34" | Car | 2012 | 0 | 2 |
|  | Cyprus | Cyprus | Cyp12 | N 34° 39' 5"<br>E 33° 0' 17" | Car | 2012 | 0 | 1 |
|  |  | Cyprus | NA | N 34° 39' 5"<br>E 33° 0' 17" | Car | 2013 | 0 | 0 |
| Captive | France | Nice | CapNiceA | N 43° 41' 23"<br>E 7° 18' 6" | Car | 2013* (2011) | 50 | 0 |
|  |  | Nice | CapNiceB | N 43° 41' 23"<br>E 7° 18' 6" | Car | 2013** (2011) | 31 | 0 |
|  |  | Valence | CapVal | N 44° 58' 21"<br>E 4° 55' 39" | Che/Pea/Haz | 2013 (2013) | 50 | 0 |
|  | Israel | Tel-Aviv | CapIsr | Not Available | Not Available | 2013 (2011) | 50 | 0 |

\* sampled in July 2013
\*\* sampled in October 2013
†Several sites were searched in these locations but no wasps were found

**Figure 1.**
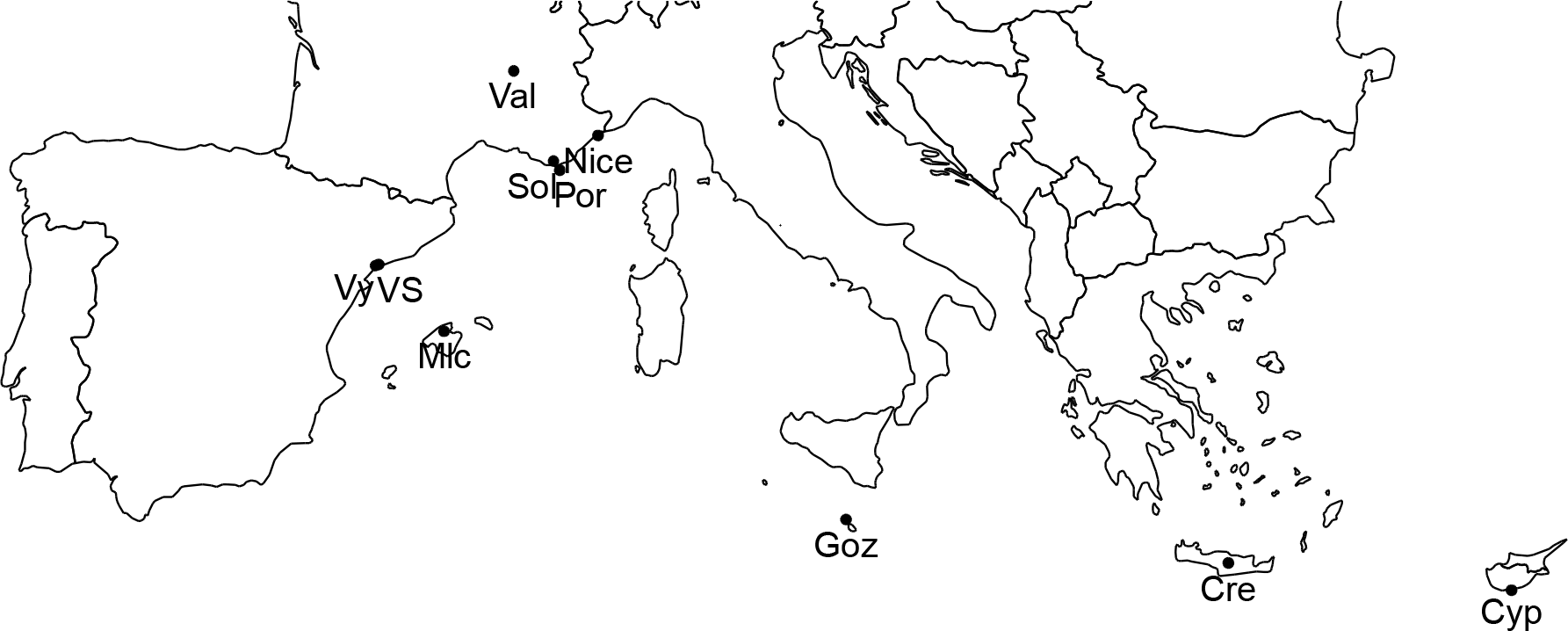
Location of field sampling. Cre, Cyp, Goz, Mlc, Nice, Sol, Val, VS and Vy are respectively acronyms for Crete, Cyrpus, Gozo, Mallorca, Nice, Solliès, Valence, Vila-Seca and Vinyols. As no wasps were found in Porquerolles, Corsica anSicily,these locations are not presented here.

In addition, males were searched in eighteen locations of the Mediterranean basin between 2011 and 2013 during late summer and autumn. Five sites were located on mainland in Nice (France, 2011 and 2013), Solliès (France, 2013), Vinyols and Vila-Seca (Spain, 2013). In 2012, we searched *V. canescens* males in five sites located on islands: Mallorca (Spain), Gozo (Malta), Crete (Greece), Sicily (Italy) and Cyprus. Several sites per island were sampled. In 2013, the same protocol was followed to capture males in French islands (Porquerolles and Corsica), in Mallorca (Spain),Cyprus and in Crete (Greece) (Table 1 and Figure 1). We selected field sites with at least 10 individual hostplants (i.e. carob, fig, pomegranate, citrus, palm or walnut trees; (Salt, 1976). On each site, ten male traps were evenly distributed. They were hung in host trees at 50 to 150 cm from the ground. Traps were checked after 48h. If one *V. canescens* individual was caught, 20 to 40 traps were added on the site. *V. canescens* malesare attractedbythe synergy of pheromones produced by females and kairomones from their hosts (Metzger et al., 2010b). Traps baited with extracts from these semiochemicals were designed to attract and capture males. A trap consisted of a 125 mm è200 mm yellow sticky sheet, in the center of which was hung a vial containing the extract. To prepare the extract,100 g of host-rearing medium containinglarvae were immerged during one hour in 200 ml of hexane. The solution was filtered andone female was soaked in 100μl for 3 hours. The female was then removed and the extract was stored at-20°C. A few hours before being used in the field, the extract was evaporated by heating the vial on a hot plate (this step was skipped for the 2013 campaign, after it was shown that evaporation occurred within a few hours in the field). As for females, traps were hung within host trees, and all *V. canescens* males that stuck on the traps were collected and preserved individually in 96% ethanol.

In parallel, we collected males in three captive (laboratory mass-reared) populations differing in their history and geographic origin (Table 1): (i) a 25-30 generation-old population founded with about 120 females from Nice, (ii) a 10-15 generation-old population founded with about 100 females from Valence, and (iii) a 15-20 generation-old population founded with 11 females from Israel. Hence, these captive populations experienced different intensities of founding effects, genetic drift, and inbreeding. The three populations were maintained with a standard protocol on larvae of the host *E. kuehniella* obtained by rearing 50 mg of *E. kuehniella* eggs (about 2000 eggs) provided by Biotop (Livron-sur-Drôme, France) in plastic boxes (8 × 12 × 25 cm) set with250 g of wheat semolina. Each week, three boxes containing 2^nd^ to 5^th^ instar host larvae were inoculated with 50 males and 50 females of *V. canescens* emerging from all available (about six) rearing boxes.

### Development of microsatellite genetic markers

Using a DNEasy Tissue Kit from Qiagen (QIAGEN, Hilden, Germany), DNA was extracted from a single 1.5 ml tube containing a pool of 20 adult females of *V. canescens* recently collected from the Nice population and kept in the laboratory for a few generations. The obtained DNA solution was enriched in microsatellites and pyrosequenced by the company Genoscreen (Lille, France), following the protocol described in Malausa *et al.* (2011). Using the iQDD program (Meglecz et al., 2010), primer pairs were designed for 675 microsatellite loci (Malausa et al., 2011), among which 124 were screened in monoplex PCR on DNA of one thelytokous and six arrhenotokous *V. canescens* females from three populations: Valence, Nice and Antibes (geographic coordinates: N 43° 33′ 51″, E 7° 7′ 28″). Six primer pairs designed by Mateo Leach *et al.* (2012) were also tested following the same protocol.

The DNA extracts used for these monoplex PCR were obtained using the DNeasy Tissue Kit of QIAGEN (Hilden, Germany). Each monoplex PCR contained 2μl of DNA, 5μl of 2X QIAGEN Multiplex PCR Master Mix and 0.2 μM of each primer. The total volume was adjusted to 10μl with ultrapure water. PCR was performed as follows: a step of denaturation at 95°C during 15 minutes, followed by 25 cycles of 30 seconds of denaturation at 94°C, 90 seconds of annealing at 58°C and 60 seconds of extension at 72°C, and a final extension step of 30 minutes at 60°C. PCR products were separated on a 2% agarose gel stained with ethidium bromide to detect amplification of DNA fragments. For 37 successfully amplified microsatellite loci, forward primers labeled with one of four fluorochroms (Applied Biosystems, Carlsbad, USA) were used in a monoplex PCR with the DNA samples and conditions previously described. Two microliters of PCR product were added to 8.75μl of Hi-Di formamide and 0.25 μl of GeneScan500 Liz size standard (Applied Biosystems). The mix was loaded on an ABI 3130XL genetic analyzer (Applied Biosystems) and alleles were scored using Gene Marker®version 1.75 (SoftGenetics, State College, USA). Monomorphic markers were discarded and the remaining markers were combined according to their size and fluorochrom in several multiplex PCR that were tested on the same seven individuals and in the same PCR conditions as previously described. Finally, we selected 19microsatellite loci amplified using two multiplex PCR (Table 2).

**Table 2.**
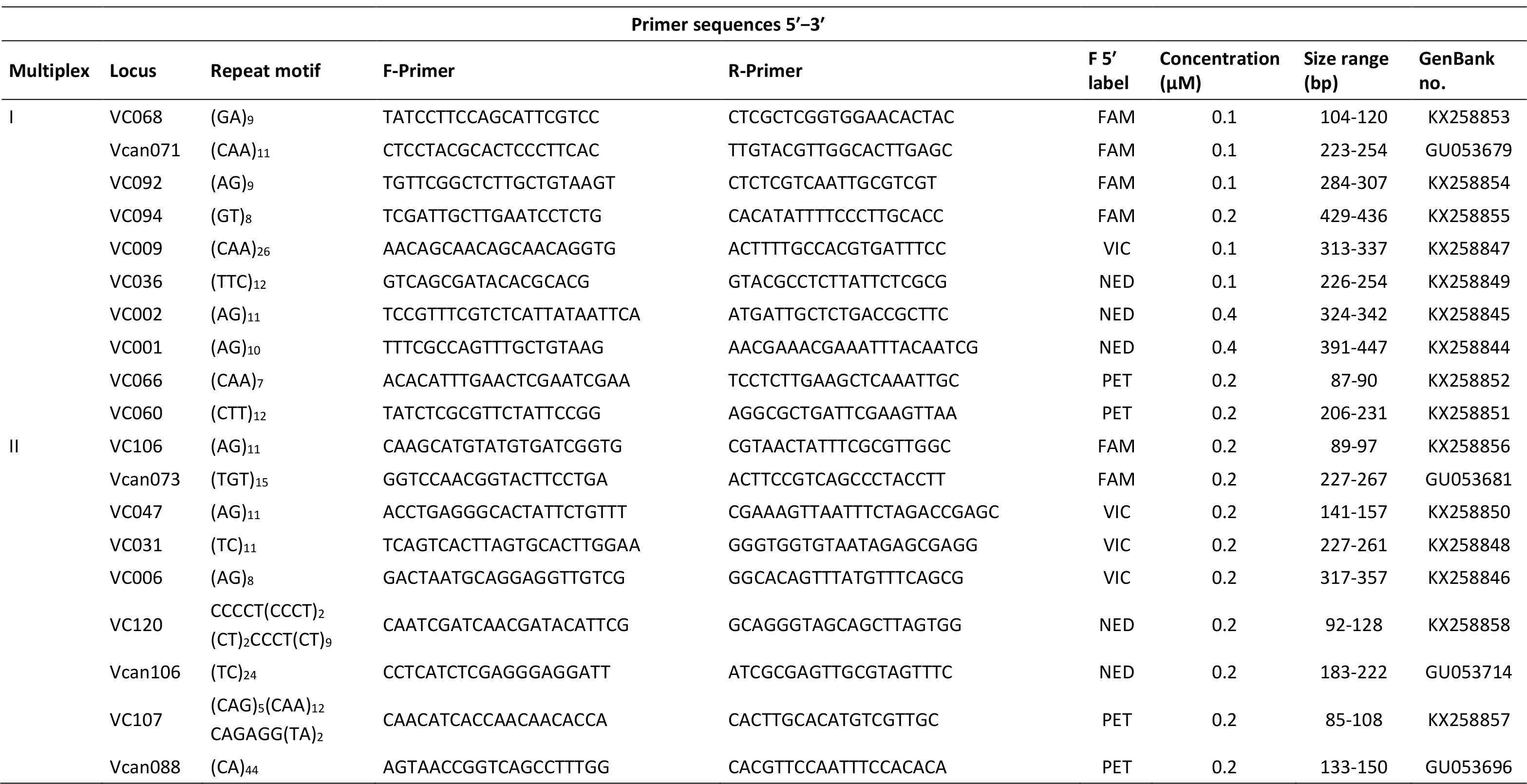
Characteristics of two multiplex PCR amplifying 19 microsatellite loci in *Venturia canescens*. All the microsatellites were developed in this study, except Vcan071, Vcan073, Vcan106 and Vcan088 that comes from Mateo Leach et al. (2012).

### Assessment of male ploidy

We used the genetic markers from the multiplex PCR I (Table 2) to detect diploid males: if a male is found heterozygous for at least one locus, it is considered diploid (Armitage et al., 2010; Souza et al., 2010; Zhou et al., 2006). However, if a diploid male is homozygous for all loci, it will be falsely scored haploid. To estimate the power of the developed microsatellite markers to correctly assess male ploidy, we calculated the probability that a diploid male is homozygous at all the 10 loci of the multiplex and hence,falsely considered haploid. For this, we first calculated the probability that a diploid male produced by a brother-sister pair is homozygous for all 10 loci of the multiplex PCR I (see Annex). In parallel, we compared ploidy assessed on the same individuals via microsatellite genotyping and flow cytometry. Measure of male ploidy by flow cytometry is based on DNA quantity in cell nuclei (haploid nuclei being expected to have twice less DNA than diploid nuclei). To produce enough diploid males for analysis, brother-sister pairs were formed with individuals from a captive population. In offspring, 39 males were collected and killed: the thorax was used for genetic analyses and the head for flow cytometry.

We performed flow cytometry analyses as described in de Boer *et al.* (2007). Only the head of insects was used for flow cytometry because endoreduplication doubles the quantity of DNA in cells of other parts of the body of haploid males, making haploid and diploid nuclei undistinguishable (Aron et al., 2005). To isolate cell nuclei, each insect head was crushed in 0.5 ml of Galbraith buffer by turning the B pestle 20 times ina Dounce tissue grinder maintained on ice. The homogenate was filteredthrough a 40μmcell strainer cap. Cell nuclei were stained with 20μl propidium iodide at 1 mg/ml. Nuclei were analyzed ona LSRII Fortessa flow cytometer (BD Biosciences, San Jose, USA) with an excitation wavelength of 561 nm. Using FACSDiva Version 6.1.3 (BD Biosciences),we analyzed 10 000 nuclei per sample in a region that excluded doublets and debris. Flow cytometric DNA-histograms of diploid females were used as reference to identify diploid males.

### Population genetics

#### Validation of microsatellites on female wasps

To be used in population genetics studies, each marker must have several detectable alleles, and must not be linked with any other marker or locus under selection. We therefore measured genetic polymorphism, estimated frequencies of null alleles, and test for departures from Hardy-Weinberg (HW) equilibrium and linkage disequilibrium on the two mainland populations from Valence and Nice where enough females were captured in 2010 (Table 1). DNA of each captured female was extracted with the PrepGem Insects kit (ZyGEM, Hamilton, New Zealand). DNA extraction, PCR and genotyping were performed as previously described. The GENEPOP software version 4.3 (Rousset, 2008) was used to calculate the number of alleles, the expected and observed heterozygosity, and to test for HW equilibrium and linkage disequilibrium between loci. We estimated null allele frequencies with the FreeNA program (Chapuis and Estoup, 2007).

#### Population structure and male ploidy

For all males captured, DNA was extracted, PCR were performed with multiplex I, and genotypes analyzed as previously described. Only males with at least eight loci genotyped were used for the analysis. Coexistence of male haploidy and diploidy impedes the estimation of classical population genetic statistics such as observed and expected heterozygosity or Wright´s F statistics. We therefore computed a value of allelic richness per locus and per population with FSTAT version 2.9.3.2 (Goudet, 1995), and calculated the number of private alleles in each population (*i.e.* the number of alleles specific to a population), because allelic richness can be standardized independently of sample size and both measures are independent of heterozygosity. The allelic richness was computed as in El (Mousadik and Petit, 1996), *i.e.* a mean number of alleles for each locus in each population, with populations weighted in inverse proportion to their sample sizes in order to give the same weight to all populations. As FSTAT is not able to handle haploid and diploid data in the same analysis, we merged the haploid data from pairs of males to create “false diploids” without changing allelic frequencies. We then analyzed all “diploids” in a single run.

The ploidy of each captured male was deduced from its genotype at microsatellite markers from multiplex PCR I. A male was considered diploid if it was heterozygous at one or more locus.

### Statistical analysis

We used a hierarchical generalized linear model (binomial error and logit link) to assess the effect of population structure on the proportion of diploid males (DMP). Population structure was characterized by different nested explanatory variables: *Allelic richness* (All̲rich) and *number of private alleles* (Priv̲all) nested inthe *habitat* type (Habit), *i.e.* Mainland, Island or Captive (see Table 1). Data were all analyzed with the statistical software R (version 3.2.2 (R Core Team, 2013). The significance of terms in the statistical model was assessed with type II sums-of-squares tested with analyses of deviance with likelihood-ratio test based on the χ^2^ distribution (*car* package (Fox and Weisberg, 2010). We used non-parametric analysis to test for population differentiation (Kruskal-Wallis test on allelic richness and the number of private alleles). Due to the low sample size (11 when using populations and three when testing the effect of habitat types) and therefore the lack of statistical power, we estimated the effect size by computing the Cohen′s *d* (Cohen, 1977; Nakagawa and Cuthill, 2007) with *effsize* library in R (Torchiano, 2015), and used the threshold provided in (Cohen, 1992) to assess the magnitude of the effect, *i.e.* |*d*| <0.2:negligible, |*d*| <0.5: small, |*d*|<0.8: medium, otherwise large.

In all multiple analyses such as tests for Hardy-Weinberg equilibrium, p-values were corrected for multiple testing via the False Discovery Rate (FDR) procedure (Benjamini and Hochberg, 1995). Means are thereafter presented with standard errors unless indicated otherwise.

## Results

### Insect sampling and microsatellite markers

108 females were collected between 2010 and 2013 and 627 males were collected in 11 populations (Table 1 and Figure 1). No males were found in the French, Italian, Greek, Malta and Cyprus islands and, as only one to three females were sampled in these locations, they were not used for the microsatellite development. Therefore, we used 75 females from Nice and Valence (2010) for the development of genetic markers.

#### Validation of microsatellite markers for population genetic studies

For all the nineteen microsatellites developed, amplification was obtained for all the arrhenotokous *V. canescens* females trapped in Nice (Nice10 population) and in Valence (Val10 populations). The number of alleles ranged from 2 to 14 (mean number of alleles=7.95 ± 0.76, Table 3). The two multiplex PCR displayed a similar range (Multiplex I: 2-12 alleles, Multiplex II: 4-14 alleles) and mean number of alleles (Multiplex I: 7.3 ± 0.92, Multiplex II: 8.67 ± 1.25, t-test: *t*=−0.88, *df*=15, *p*=0.39). The markers had a maximum frequency of null alleles equal to 8.5% (Vcan073 locus, Val10 population; mean: 1.7% ± 0.6 in Nice10 population and 1.8% ± 0.7 in Val10 population) and a null median frequency for both populations. No departure from the HW equilibrium was found in any of the populations (Table 3), and the observed heterozygosity (Ho) was similar in the two multiplex PCR in both populations (Mean Ho Multiplex I: 0.63 ± 0.03, Multiplex II: 0.66 ± 0.03; t-test: *t*=−0.54, *df*=36, *p*=0.59; Table 3). Two pairs of microsatellites were detected as linked when both populations were considered: Vcan073/Vcan106 (χ^2^=∞, *df*=4, *p*<0.0001) and VC107/Vcan106 (χ^2^=24.2, *df*=4, *p*=0.0063). However, when populations were analyzed separately, the pairs of loci were linked only in one population (Vcan073/Vcan106 in Val10, and VC107/Vcan106 in Nice 10), we therefore decided to keep all loci. Therefore, both multiplex PCR can be used for genetic studies with equivalent performance.

**Table 3.** Genetic diversity at 19 microsatellite loci for natural populations of *Venturia canescens* sampled in 2010 near Nice and Valence, South East of France. *n*: numberof females analysed; H_E_: expected heterozygosity; H_o_: observed heterozygosity; HW P-value: P-value of the test for Hardy-Weinberg equilibrium after FDR correction.

| Multiplex | Primer name | Number of alleles | Population |  |  |  |  |  |  |  |
| --- | --- | --- | --- | --- | --- | --- | --- | --- | --- | --- |
|  |  |  | Nice10 (*n* = 44) |  |  |  | Val10 (*n* = 37) |  |  |  |
|  |  |  | *H<sub>E</sub>* | *H<sub>O</sub>* | HW P-value | Null allele frequency | *H<sub>E</sub>* | *H<sub>O</sub>* | HW P-value | Null allele frequency |
| I | VC068 | 7 | 0.742 | 0.727 | 0.848 | 0.012 | 0.826 | 0.865 | 0.771 | 0.000 |
|  | Vcan071 | 11 | 0.809 | 0.750 | 0.493 | 0.036 | 0.798 | 0.838 | 0.728 | 0.000 |
|  | VC092 | 7 | 0.636 | 0.705 | 0.390 | 0.000 | 0.600 | 0.676 | 1.000 | 0.000 |
|  | VC094 | 4 | 0.592 | 0.545 | 0.771 | 0.034 | 0.511 | 0.541 | 0.813 | 0.000 |
|  | VC009 | 8 | 0.697 | 0.864 | 0.135 | 0.000 | 0.649 | 0.703 | 1.000 | 0.000 |
|  | VC036 | 7 | 0.540 | 0.568 | 0.771 | 0.000 | 0.450 | 0.432 | 0.771 | 0.000 |
|  | VC002 | 7 | 0.577 | 0.591 | 0.813 | 0.026 | 0.716 | 0.757 | 0.771 | 0.000 |
|  | VC001 | 12 | 0.600 | 0.477 | 0.370 | 0.090 | 0.655 | 0.595 | 0.176 | 0.046 |
|  | VC066 | 2 | 0.487 | 0.386 | 0.509 | 0.069 | 0.491 | 0.486 | 1.000 | 0.003 |
|  | VC060 | 8 | 0.549 | 0.591 | 0.927 | 0.000 | 0.558 | 0.568 | 0.135 | 0.011 |
| II | VC106 | 4 | 0.711 | 0.750 | 0.976 | 0.000 | 0.728 | 0.676 | 0.307 | 0.019 |
|  | Vcan73 | 10 | 0.672 | 0.818 | 0.746 | 0.000 | 0.706 | 0.541 | 0.135 | 0.092 |
|  | VC047 | 5 | 0.573 | 0.636 | 0.821 | 0.000 | 0.607 | 0.541 | 0.396 | 0.013 |
|  | VC031 | 14 | 0.650 | 0.591 | 0.390 | 0.048 | 0.674 | 0.595 | 0.493 | 0.083 |
|  | VC006 | 11 | 0.673 | 0.773 | 0.813 | 0.000 | 0.593 | 0.595 | 0.771 | 0.000 |
|  | VC120 | 7 | 0.515 | 0.500 | 0.771 | 0.000 | 0.593 | 0.459 | 0.135 | 0.059 |
|  | Vcan106 | 14 | 0.875 | 0.886 | 0.400 | 0.008 | 0.836 | 0.919 | 0.746 | 0.000 |
|  | VC107 | 6 | 0.724 | 0.705 | 0.493 | 0.000 | 0.603 | 0.622 | 0.479 | 0.022 |
|  | Vcan088 | 7 | 0.644 | 0.682 | 0.813 | 0.009 | 0.489 | 0.541 | 0.821 | 0.000 |

#### Validation of markers for ploidy assessment

The probability that a diploid male produced by sib-mating is homozygous for all microsatellite loci of the multiplex I was low enough (Prob.=0.0023) to rely on microsatellites for ploidy assessment. This was confirmed by the congruence between flow cytometry and microsatellite genotyping results. Haploid and diploid males were easily discriminated by flow cytometry, with diploid males presenting a profile similar to that of diploid females (Fig. 2). For the 39 males tested, the ploidy measured by flowcytometry and genotyping analyses matched perfectly.

**Figure 2.**
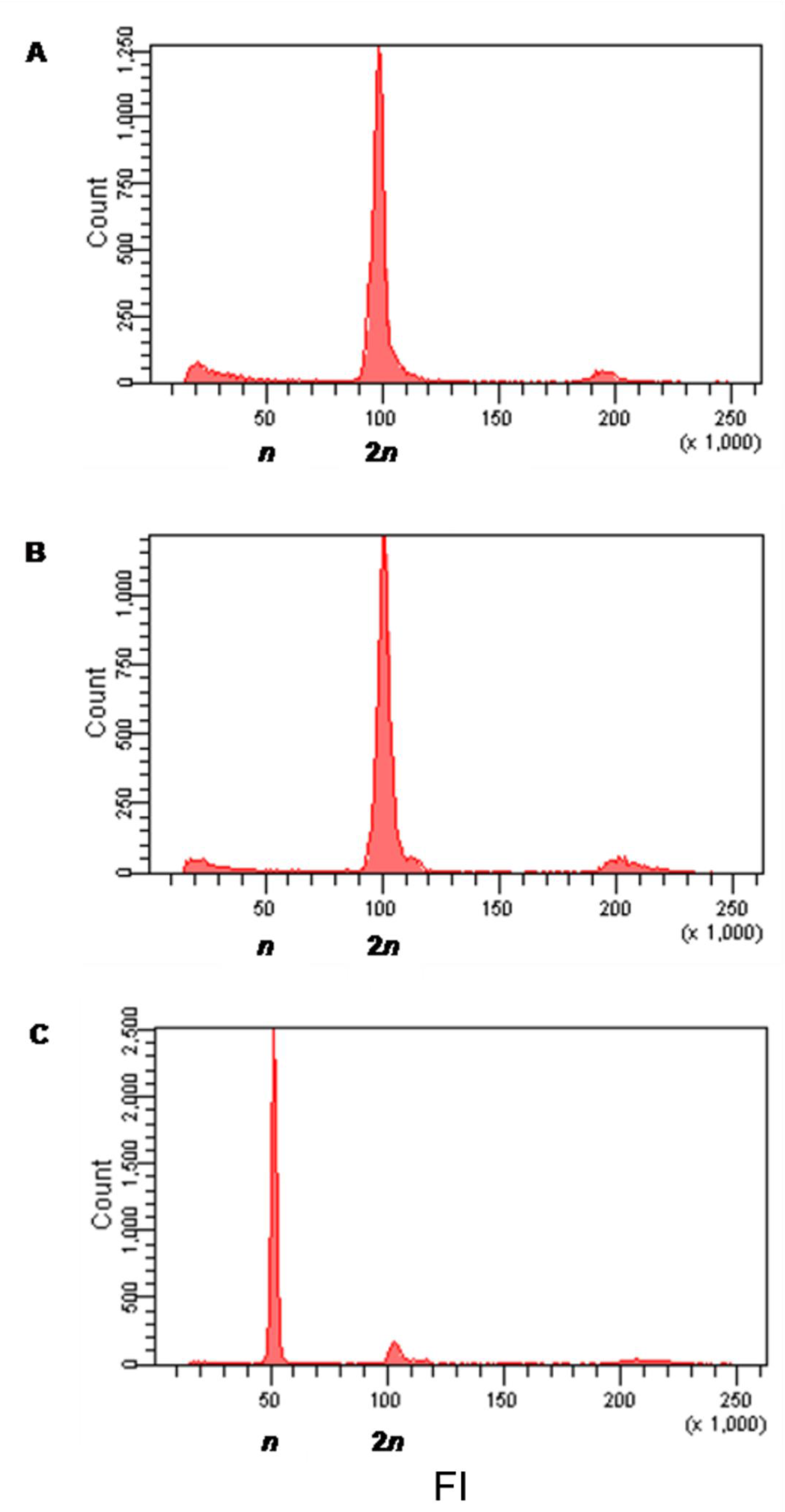
Flow cytometric histograms of the number of nuclei registered as a function of their fluorescence intensity (F1),for a representative female (A), diploid male(B) and haploid male (C). F1 is expressed in an arbitrary unit calibrated to value 100atthe fluorescence intensity with the highest number of nuclei registered in females,which are known to be diploid.

### Population Genetic Structure and Proportion of Diploid Males

After genotyping with the markers of multiplex I, we discarded 28 males for which more than one loci did not amplified. As a result, 599 males (95.5%) were successfully genotyped and were used in the analysis.

#### Population differentiation

The population structure was estimated through different parameters such as allelicrichness (All̲Rich) and the number of private alleles (Priv̲all) from male genotyping data (Table 4). The number of private alleles was marginally significantly different between habitat types (Mainland,Island and Captive, Kruskal-Wallis test, χ^2^=5.68, *df*=2, *p*=0.058) and allelic richness significantly differed between habitat types (Kruskal-Wallis test, χ^2^=6.72, *df*=2, *p*=0.035) and populations (Kruskal-Wallis test, χ^2^=18.51, *df*=10, *p*=0.047). Both measures shared the same trend suggesting that the diversity is the lowest in captive populations and the highest in mainland (Table 4). Effect sizes were medium or large for each comparison for both variables (Mainland-Island, All̲Rich: *d*=2.35, Priv̲all: *d*=1.09; Mainland-Captive, All̲Rich: *d*=1.56, Priv̲all: *d*=1.52;Island-Captive, All̲Rich: *d*=0.67, Priv̲all: *d*=1.41). Allelic richness ofthe captive population from Israel (CapIsr) appeared different from all the other populations except the captive populations from Nice and Valence (CapNiceB and CapVal, pairwise t-tests between populations with FDR correction for multiple testing, data not shown). As the CapIsr population was highly differentiated from the other populations, it could potentially drive the significant trend observed. Indeed, when allelic richness was compared without CapIsr, the difference between populations disappeared (Kruskal-Wallis test, χ^2^=4.745, *df*=9, *p*=0.856) and the difference between habitat types became only marginally significant (Kruskal-Wallis test, χ^2^=5.804, *df*=2, *p*=0.055).

**Table 4.** Characteristics of *Venturia canescens* populations based on the analysis of males. Allelic richness and mean number of alleles were computed with FSTAT software. This software being unable to handle both haploid and diploid data in the same analysis, the co-occurrence of haploid and diploid males constrained us to merge pairs of haploid data to create “false diploid” males. The results presented are mean ± SE.

| Location | Population | Number of sampled males | Number of genotyped haploid males | Number of genotyped diploid males | Mean number of alleles | Overall allelic richness | Number of private alleles |
| --- | --- | --- | --- | --- | --- | --- | --- |
| **Mainland** |  |  |  |  |  |  |  |
| France | Nice11 | 190 | 172 | 10 | 6.78 ± 0.92 | 3.37 | 8 |
| | Nice13 | 90 | 85 | 2 | 6.00 ± 0.75 | 3.52 | 0 |
| | Sol13 | 16 | 12 | 1 | 3.78 ± 0.43 | 3.58 | 2 |
| Spain | VS13 | 58 | 49 | 3 | 5.67 ± 0.78 | 3.41 | 3 |
| | Vy13 | 33 | 30 | 1 | 5.22 ± 0.64 | 3.68 | 4 |
| *Mean Mainland* | | 77.4 ± 3.5 | 69.6 ± 3.4 | 3.4 ± 0.9 | 5.49 ± 0.34 | 3.51 ± 0.14 | 3.4 |
| **Island** |  |  |  |  |  |  |  |
| Spain | Mlc12 | 21 | 18 | 3 | 4.22 ± 0.43 | 3.27 | 0 |
| | Mlc13 | 38 | 33 | 4 | 4.56 ± 0.47 | 3.25 | 1 |
| *Mean Island* | | 29.5 ± 1.6 | 25.5 ± 1.5 | 3.5 ± 0.3 | 4.39 ± 0.31 | 3.26 ± 0.18 | 0.5 |
| **Captive** |  |  |  |  |  |  |  |
| France | CapNiceA | 50 | 43 | 7 | 4.33 ± 0.37 | 3.32 | 0 |
| | CapNiceB | 31 | 26 | 1 | 3.78 ± 0.52 | 3.04 | 0 |
| | CapVal | 50 | 48 | 1 | 3.67 ± 0.44 | 3.08 | 0 |
| Israël | CapIsr | 50 | 42 | 8 | 2.44 ± 0.24 | 2.05 | 0 |
| *Mean Captive* | | 45.3 ± 0.7 | 39.8 ± 0.8 | 4.3 ± 0.9 | 3.56 ± 0.23 | 2.87 ± 0.15 | 0 |

#### Diploid male proportion (DMP) and genetic diversity

Diploid males were found in all populations, but in variable proportions (from DMP=0.02 ± 0.02 in CapVal to DMP=0.16 ± 0.05 in CapIsr; Table 4 and Figure 3). As expected, the DMP was affected by habitat type, with a negative relationship between the DMP and the populations' habitat types (LR χ^2^=7.07,*df*=2, *p*=0.029, Table 5 and Figure 4). When removing the CapIsr population, the effect of the habitat type became non-significant (*p*=0.105) but we still found a negative influence of allelic richness (nested within habitat types) on DMP (LR χ^2^ =6.80, *df*=2, *p*=0.033, Table 5).

**Figure 3.**
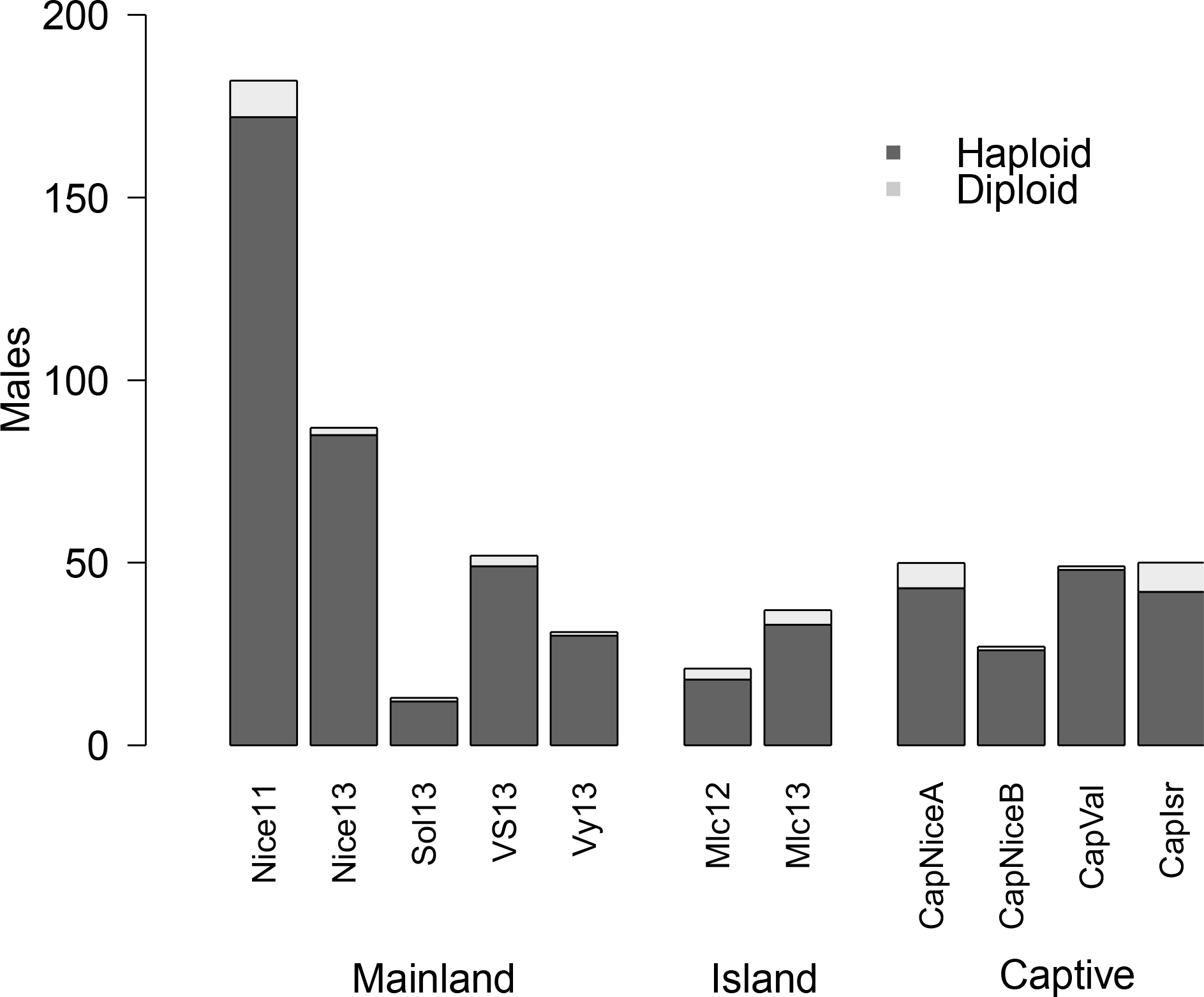
Number of haploid and diploid males in the 11 populations within each habitat type.

**Table 5.**
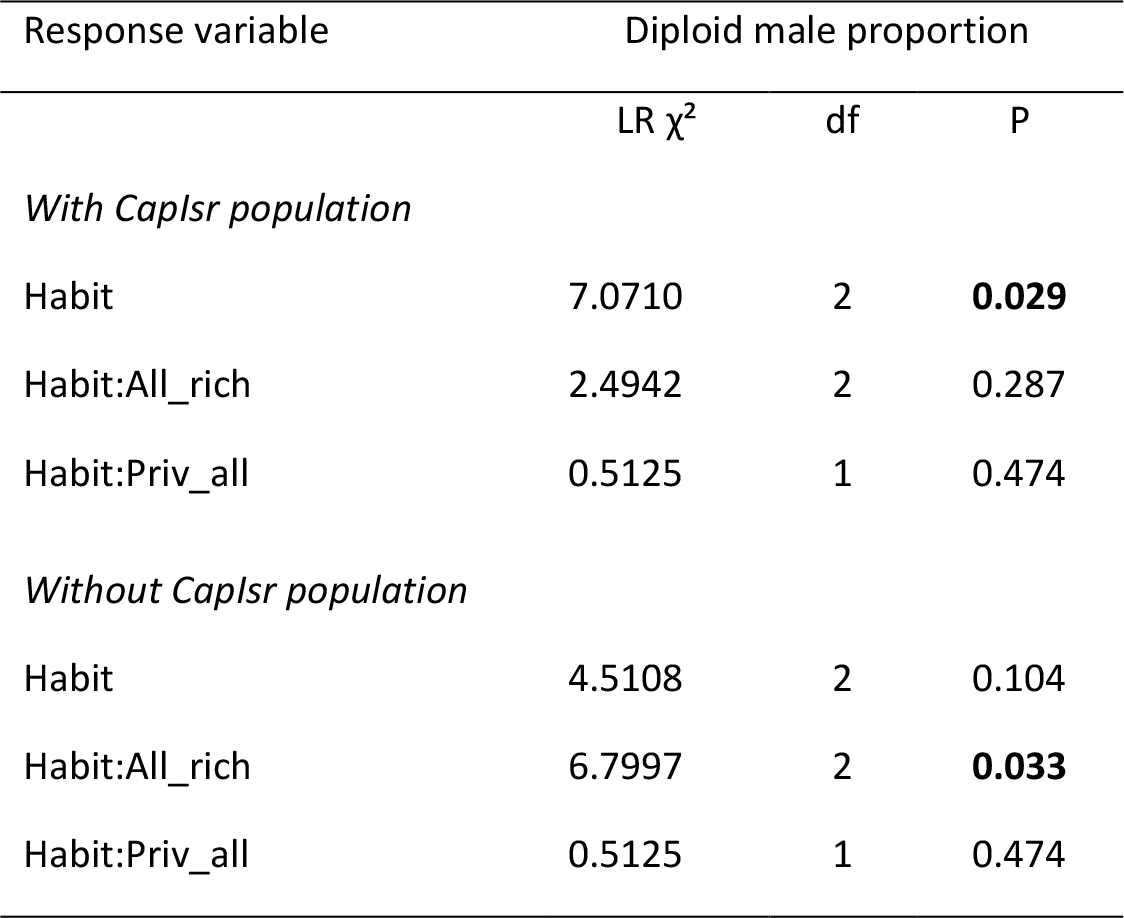
Effect of habitat type and genetic diversity on diploid male proportion. LR, Likelihood ratio; df, degrees of freedom; Habit, habitat type; All̲rich, allelic richness; Priv̲all, private alleles. *P*<0.05 highlighted in bold.

**Figure 4.**
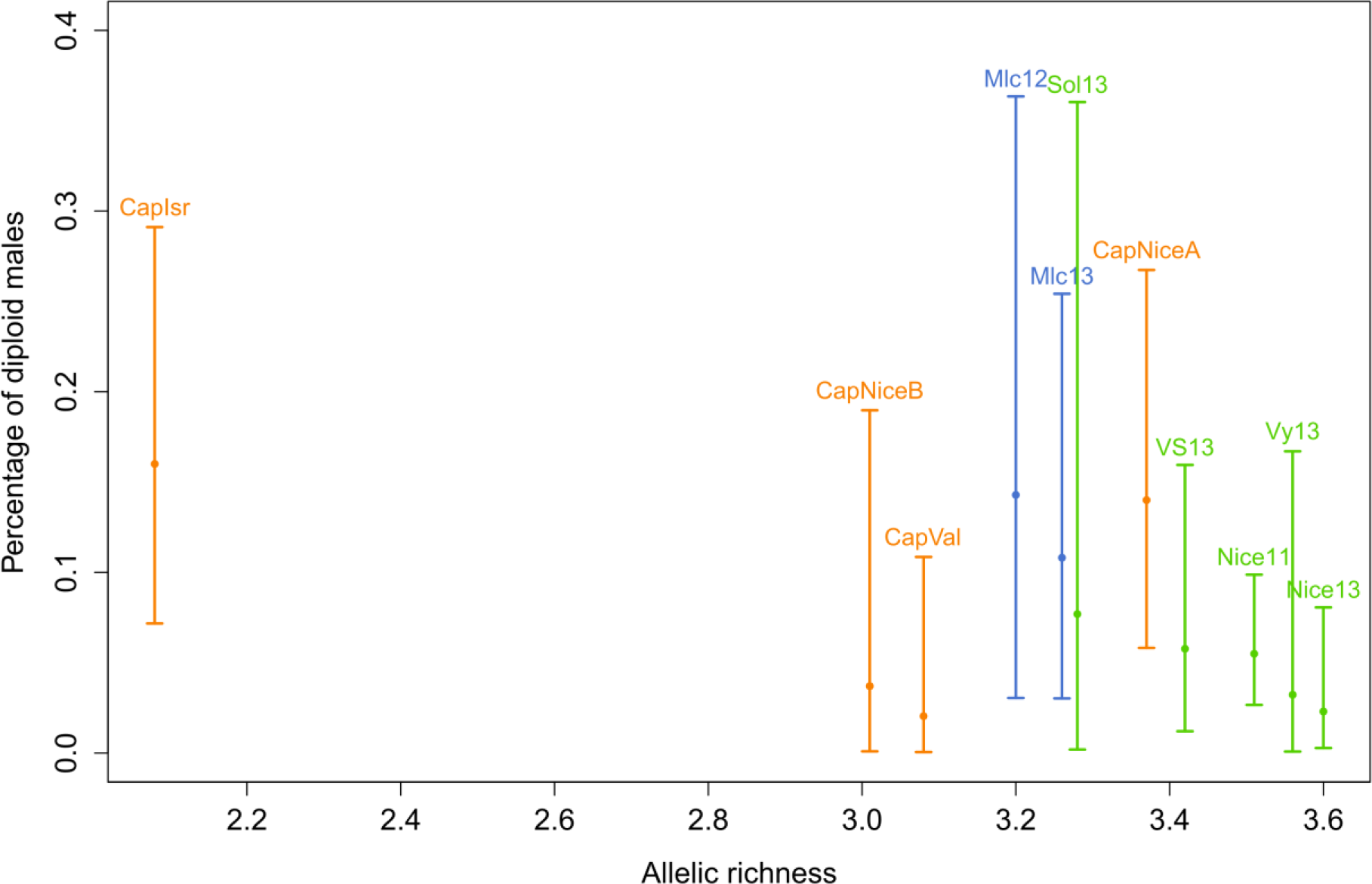
Percentage of diploid males according to allelic richness in the 11 populations, with 95% confidence intervals. Colours represent the three habitat types of locations: light green for the mainland populations, light blue for the island populations and orange for the captive populations.

## Discussion

We developed highly variable microsatellite markers for the parasitoid wasp Venturia canescens. Polymorphism, Hardy-Weinberg equilibrium, absence of linkage disequilibrium and negligible null allele frequencies make these markers suitable for population genetic studies. In addition, relatively high rates of heterozygosity enable a reliable measure of male ploidy, confirmed by flow cytometry. Using these markers, we estimated within-population genetic diversity and the proportion of diploid males in eleven locations around the Mediterranean Sea, as well as in captive populations. We found a tendency for genetic diversity to be reduced in small and isolated habitats, and the proportion of diploid males to increase with decreasing genetic diversity. These patterns are expected theoretically in a scenario whereby reduced population size yields decreased genetic diversity (and decreased allelic polymorphism at the sl-CSD locus), increased proportion of diploid males, decreased population growth rate and, if dramatic enough, population extinction (a scenario referred to as the diploid male vortex; Zayed and Packer 2005). Hence, our results suggest that habitat type and population history are key features that should be considered when studying genetic diversity in parasitoids with sl-CSD.

The effect of habitat type on genetic diversity was marked by a lower diversity in captive populations. These bottlenecked and completely isolated populations were characterized by a lower allelic richness compared to either island or mainland populations, and a lower number of private alleles compared to mainland populations (Table 3). Because the number of private alleles is highly dependent on sampling effort, the effect of habitat type on the number of private alleles could simply result from the high number of individuals captured in the mainland population of Nice. Besides, the captive population from Israel was highly differentiated from all populations as a result of a very small foundress number (11 females). This population had the lowest allelic richness and mean number of alleles and hence, it is likely that these extreme values had driven the effect of habitat type. Indeed, the population effect disappeared when discarding the population from Israel. Nonetheless,the effect of habitat type remained marginally significant, suggesting that the trend of higher allelic richness in mainland compared to island and captive populations is indeed real.

The decrease of genetic diversity co-occurring with higher rates of inbreeding in small and isolated populations, such as island populations, has been documented in various species (mammals, birds, fishes, insect or plants, (Frankham, 1997; Furlan et al., 2012). In *Venturia canescens*, the higher genetic diversity in mainland compared to captive populations could be explained by efficient dispersal. In this species, adults are good dispersers (Desouhant et al., 2003), a feature that corroborates the absence of genetic structure at the scale of Southeastern France (Schneider et al., 2002). In island and captive populations, founder effect and genetic drift combined with constrained dispersal should lead to genetic erosion, a phenomenon that is also observed in other species of the order Hymenoptera such as bumblebees or orchid bees (Boff et al., 2014; Ellis et al., 2006; Schmid-Hempel et al., 2007).

Consistently with these effects of habitat type on diversity at microsatellite loci, the proportion of diploid males was affected by the habitat type with more frequent diploid males in island and captive populations than in mainland locations. The highest proportion (16%) was observed in the captive population from Israel, which is congruent with the lowest genetic diversity estimated. In other species, diploid male production was also found higher in isolated population compared to larger genetically-diverse populations (Kukuk and May, 1990). Overall, the proportion of diploid males in *V. canescens* (6.8% overall, CI 95% [5.0%-9.2%]) is in the order of magnitude of estimations in other species of parasitoids with sl-CSD: 10% in a native population of *Cotesia glomerata* (Ruf et al., 2013) and 15% in a population of Cotesia rubecula introduced for biological control (de Boer et al., 2012).

Behaviours such as natal dispersal and mate choice can mitigate the effects of decreased genetic diversity and hence attenuate diploid male production. In *C. glomerata*, a biased fertilization occurs and limits genetic incompatibility, leading to a DMP lower than expected in field populations (Ruf et al., 2013). On the contrary, the lack of mate discrimination in *C. rubecula* could explain the relatively high DMP observed in its field populations (de Boer et al., 2012). Laboratory studies in Bracon brevicornis have reported an avoidance of mating with a partner harboring the same allele at the sl-CSD locus (*i.e.* avoidance of matched matings; Thiel et al., 2013), and sib-mating avoidance was observed in *V. canescens* (Metzger, Bernstein, et al. 2010). Such behaviours should reduce the DMP and thus lessens the production of unfit offspring (Chuine et al., 2015; Parker, 1983).

The higher proportion of diploid males in captive and island populations of *Venturia canescens* could result from a lower genetic diversity in isolated populations, or a downgrade in discrimination of females against related males. In a number of species, females adapt future mate choice according to the genotype of their first mate (lizards: (Breedveld and Fitze, 2015; Laloi et al., 2011), beetles: (Dowling et al., 2007), Drosophila: (Chapman and Partridge, 1998)). This hypothesis may not hold for *V. canescens* because females are monoandrous (Metzger et al., 2010b). However,as island or captive populations are genetically eroded, the probability for a female to encounter a male bearing a similar allele atthe *csd* locus is higher; possibly, successive encounters with low-quality males before mating could lead toa strategic decline in sib-avoidance. This couldexplain the increasein diploid male production in small or isolated populations. At last, the presence of diploid males even in mainland populations where the genetic diversity is high and where no departure from Hardy-Weinberg equilibrium was detected could also be due to imperfect sib-mating avoidance, as observed in laboratory conditions (Metzger et al. 2010).

Our study suggests that diploid male production in insects of the order Hymenoptera reflects fitness decline resulting from inbreeding in small and isolated habitats. Hence, our study raises the question of population conservation (Zayed, 2009) and highlights that small organisms such as insects may also suffer from habitat destruction and fragmentation.

## Acknowledgements

We thank Virginie Dos Santos and Anna Chuine for their comments on the draft, Didier Crochard, Bastien Quaglietti, Virginie Dos Santos for field sampling. This work was supported by the Agence Nationale de la Recherche (Sextinction project ANR-2010-BLAN-1717) and the Fédération de Recherche sur la Biodiversité (VORTEX project APP-IN-2009-052), both coordinated by XF.

## Annex

Calculation of the probability that a F2 male, produced by sibmating, is homozygous for all 10 markers from multiplex I.

A sample of 30 females was collected in a 3 to 5 generation-old captive population (founded in autumn 2009 with about 40 females from Nice). These females constitute theF0 generation and had presumably mated randomly in the population. Their offspring (generation F1) were constrained to mate between siblings (Figure A). We computed the probability that an F2 male, produced via sibmating, is homozygous for all genetic markers, using allelic frequencies and inbreeding coefficient (Fis) of F0 females.

**Figure A.**
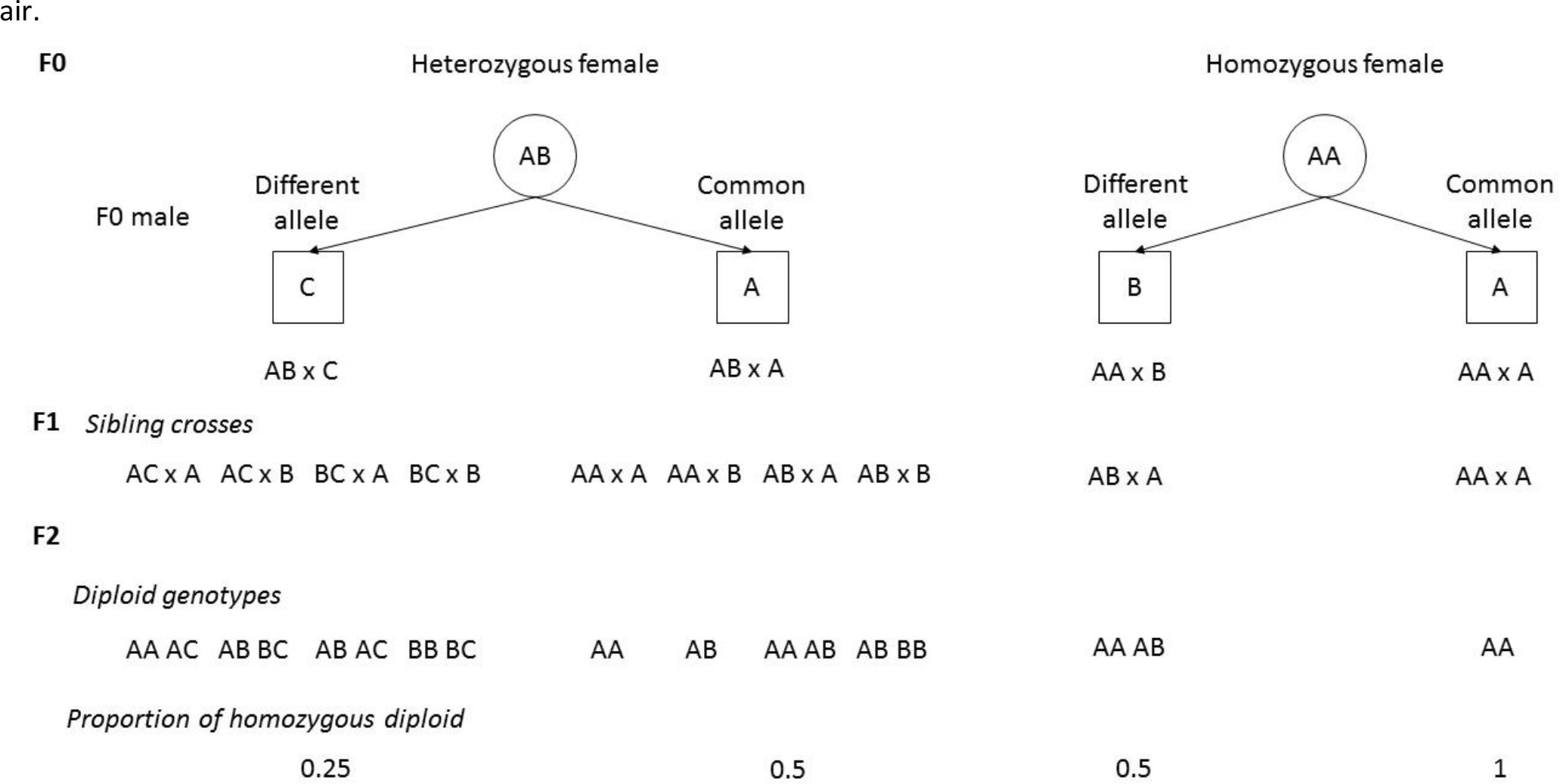
Proportion of heterozygous diploid F2 at one microsatellite locus, according to the genotype of the F0 pair

We first calculated the probability that an F2 diploid individual is homozygous at one microsatellite locus, given the genotypes of its grand-parents (F0 generation) at this locus (Figure A). Then, for each locus *k*, we calculated the probability that an F0 female is heterozygous at this locus as P*k* (Het_F0_)=(1-Fis)×He and the probability that she is homozygous as P*k*(Hom_F0_)=1-P_k_(Het_F0_), where He is expected heterozygosity at loci *k* under HW equilibrium. We calculated as follows the conditional probabilities that the F0 male carries at loci k a different (DA) or common (CA) allele with the F0 female, knowing that the female is heterozygous (Het_F0_) or homozygous (Hom_F0_):

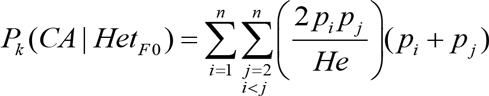

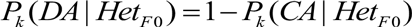

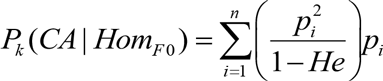

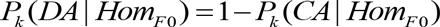

p_i_ and p_j_ are the frequencies of alleles *i* and *j*, respectively. Using the proportions of F2 homozygous from Figure A, the probability that an F2 diploid is homozygous as locus *k* was:

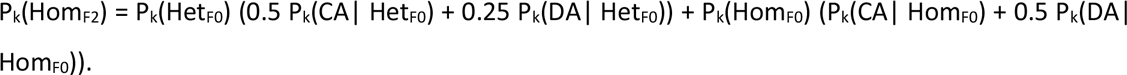

The probability that a diploid F2 is homozygous at all 10 loci from multiplex I was:

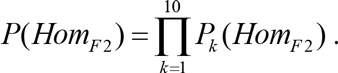

